# ABHD11 regulates 2-oxoglutarate abundance by protecting mitochondrial lipoylated proteins from lipid peroxidation damage

**DOI:** 10.1101/2020.04.18.048082

**Authors:** Peter S J Bailey, Brian M Ortmann, Jack W Houghton, Ana S H Costa, Robin Antrobus, Christian Frezza, James A Nathan

## Abstract

2-oxoglutarate (2-OG or α-ketoglutarate) relates mitochondrial metabolism to cell function by modulating the activity of 2-OG dependent dioxygenases (2-OG DDs) involved in the hypoxia response and DNA/histone modifications. However, metabolic pathways that regulate these oxygen and 2-OG sensitive enzymes remain poorly understood. Here, using CRISPR Cas9 genome-wide mutagenesis to screen for genetic determinants of 2-OG levels, we uncover a redox sensitive mitochondrial lipoylation pathway, dependent on the mitochondrial hydrolase ABHD11, that signals changes in mitochondrial 2-OG metabolism to 2-OG DD function. ABHD11 loss or inhibition drives a rapid increase in 2-OG levels by impairing lipoylation of the 2-OG dehydrogenase complex (OGDHc) – the rate limiting step for mitochondrial 2-OG metabolism. Rather than facilitating lipoate conjugation, ABHD11 protects the catalytic lipoyl domain from lipid peroxidation products formed by oxidative damage, demonstrating a requirement for a lipoyl repair pathway in human cells, and highlighting how the redox sensitivity of lipoylation modulates 2-OG metabolism.

---

The ability to sense and respond to nutrient abundance is a fundamental requirement for cell survival, and to achieve this, cells have evolved several strategies that link metabolic function to transcriptional adaptation. One such strategy is the coupling of 2-OG metabolism to gene transcription, whereby 2-OG, a key component of TCA cycle, can facilitate cell function by modulating the activity of 2-OG DDs involved in the Hypoxia Inducible Factor (HIF) response, DNA methylation, and histone modifications^1^.

The relevance of 2-OG in modulating the activity of these dioxygenases is exemplified by changes in the relative abundance of cellular 2-OG. An increased 2-OG/succinate ratio promotes embryonic stem cell pluripotency^2^, and antagonises the growth of solid organ tumours^3^ through increased hydroxymethylation of DNA (5hmC) and histone demethylation. Conversely, elevated cellular 2-OG can drive its own reduction to L-2-hydroxyglutarate (L-2-HG), which counterintuitively inhibits 2-OG DDs, leading to decreased DNA hydroxymethylation and histone demethylation, activation of the HIF response, altered T cell fate, and haematopoietic cell differentiation^4–9^. Consequently, understanding how 2-OG metabolism is regulated has broad biological implications.

Central to maintaining cellular 2-OG homeostasis is the 2-oxoglutarate dehydrogenase complex (OGDHc, also known as the α-ketoglutarate dehydrogenase complex), the rate-limiting enzyme within the TCA cycle that oxidatively decarboxylates 2-oxoglutarate to succinyl-CoA. This evolutionarily conserved enzyme also requires lipoic acid, a redox sensitive cofactor that is synthesised within the mitochondria and conjugated to a single lysine within the OGDHc E2 subunit, dihydrolipoamide S-succinyltransferase (DLST)^10–12^. The cyclical reduction and oxidation of the two thiols of conjugated lipoic acid (lipoamide to dihydrolipoamide) serves as a redox intermediate, coupling the formation of succinyl CoA to generation of NADH. The importance of DLST and its lipoylation is highlighted by the recent identification of genetic mutations leading to human disease. Patients with germline mutations in lipoic acid synthesis genes develop a severe variant of the neurological condition, Leigh syndrome^13^, and loss of heterozygosity mutations in the OGDHc lead to angiogenic tumours (pheochromocytomas and paragangliomas), similar to other hereditary cancer syndromes activating the HIF pathway^14^. However, how OGDHc function and 2-OG abundance is regulated is unclear.

Here, we use the sensitivity of the HIF pathway to 2-OG abundance to gain new insights into how 2-OG metabolism is controlled. Using genome-wide CRISPR/Cas9 mutagenesis screens, we identify an uncharacterised protein, αβ-hydrolase domain-containing 11 (ABHD11), as a mitochondrial enzyme that impairs OGDHc activity when depleted or inhibited. ABHD11 loss leads to the accumulation of 2-OG and formation of L-2-HG, which inhibits 2-OG DDs involved in the HIF response and DNA hydroxymethylation, similarly to genetic disruption of the OGDHc. ABHD11 also associates with the OGDHc and is required for catalytic activity and TCA cycle function. However, ABHD11 does not alter the constituent levels of the OGDHc. Instead, ABHD11 maintains functional lipoylation of OGDHc, protecting the enzyme from lipid peroxidation products that are propagated within mitochondria membrane following oxidative damage. Together, these studies identify a key role for ABHD11 in 2-OG metabolism, and demonstrate that OGDHc lipoylation provides a previously unappreciated mechanism for mediating an adaptive transcriptional response to changes in the mitochondrial redox environment.

## Results

### CRISPR/Cas9 mutagenesis screens identify ABHD11 as a mediator of 2-oxoglutarate dependent dioxygenase activity in aerobic conditions

To find genes involved in 2-OG metabolism we utilised the sensitivity of the HIF response to 2-OG availability, and carried out CRISPR/Cas9 mutagenesis screens in human cells using a fluorescent HIF reporter we developed^4,15^. This reporter encodes the consensus HIF responsive element (HRE) in triplicate that drives the expression of GFP fused to the oxygen and 2-OG sensitive region of HIF-1α (**Figure S1a**)^4,15^. Therefore, reporter stability is dependent on 2-OG DD activity of the prolyl hydroxylases (PHDs or EGLNs)^16,17^, which was confirmed with treatment with the PHD inhibitor dimethyloxalylglycine (DMOG), cell permeable 2-OG (dimethyl 2-OG, DM 2-OG) or incubation in 1% oxygen (**Figure S1b-d**)^4,7,15^.

Two genome-wide CRISPR sgRNA libraries were used to identify genes that when mutated activated the HIF reporter: the Brunello human genome-wide library (containing 76,441 sgRNA)^18^, and the Toronto genome-wide knockout library (containing 176,500 sgRNA)^19^. HeLa cells stably expressing the HRE-GFP^ODD^ reporter and Cas9 were transduced with each genome-wide library and iteratively sorted for GFP^HIGH^ cells by fluorescence-activated cell sorting (FACS) at day 10 and day 18 (**Figure 1a**). SgRNAs enriched by FACS were identified by Illumina HiSeq and compared to a population of mutagenized cells that had not undergone phenotypic selection (**Figure 1a, b**) (**Supplementary File S1**). All screens were conducted in aerobic conditions (21% oxygen), thereby preventing oxygen availability limiting PHD function.

**Figure 1.**
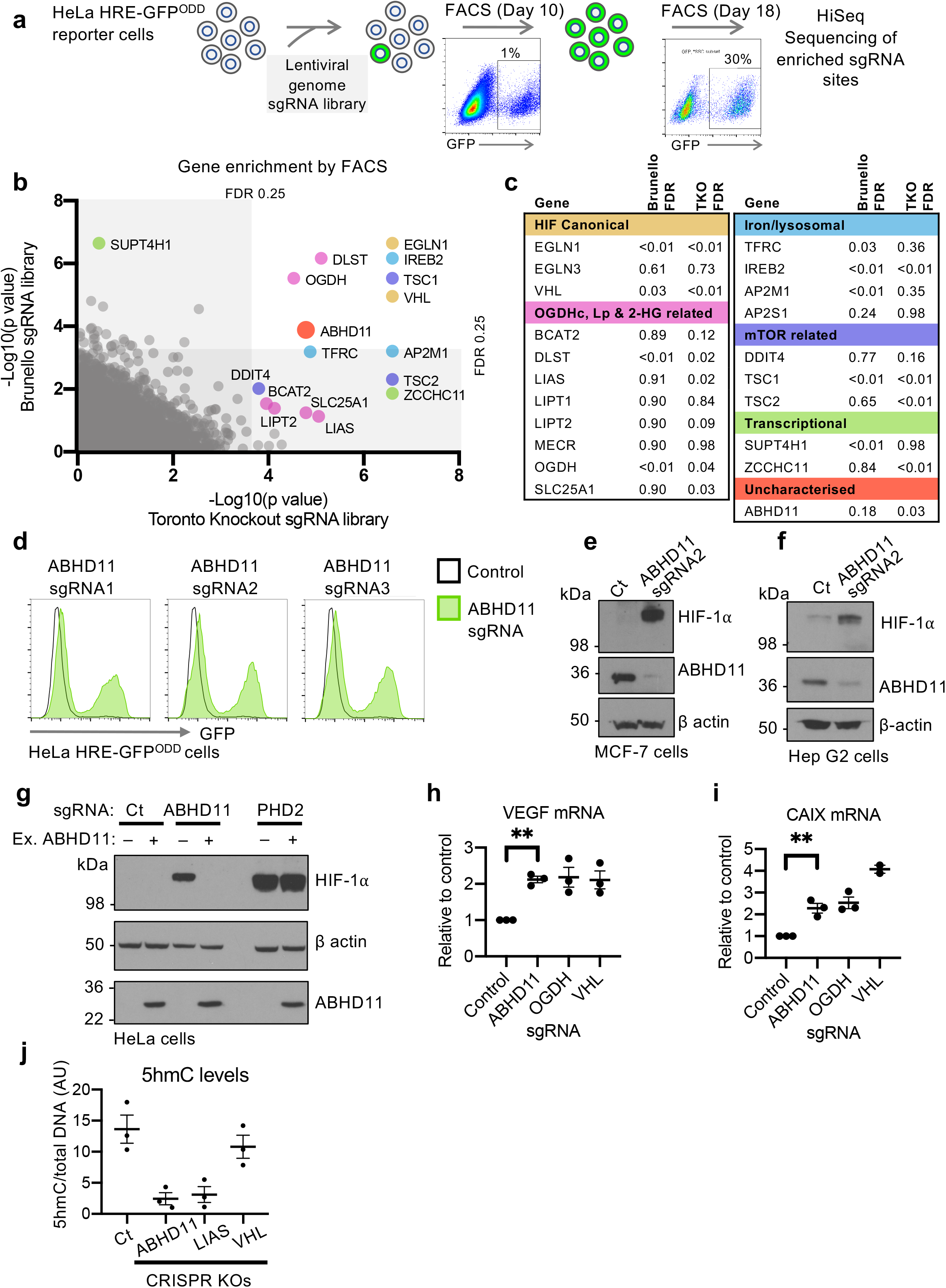
CRISPR/Cas9 mutagenesis screens identify ABHD11 as a mediator of 2-OG dependent dioxygenase activity in aerobic conditions. (**a**) Schematic of CRISPR/Cas9 mutagenesis screens. HeLa HRE-GFP^ODD^ cells were transduced and mutagenised with genome-wide sgRNA libraries (Brunello and Toronto KO). GFP^HIGH^ cells were selected by iterative FACS and sgRNA identified by Illumina HiSeq. (**b, c**) Comparative bubble plot (**b**) and table (**c**) of sgRNA enriched in the GFP^HIGH^ cells between the two genome-wide sgRNA libraries. All sgRNA enriched were also compared to a mutagenised population of HRE-GFP^ODD^ cells that had not been phenotypically selected. Genes enriched for sgRNA clustered into six main groups: (1) the canonical HIF pathway, (2) OGDHc, Lipoylation (Lp) and 2-HG related pathways, (3) intracellular iron metabolism (iron, lysosomal), (4) mTOR, (5) Transcription and (6) Uncharacterised. *EGLN1=PHD2, EGLN3=PHD3, FDR=false discovery rate.* (**d, e, f**) ABHD11 depletion results in HIF-1α accumulation. HeLa HRE-GFP^ODD^ (**d**), MCF-7 (**e**) and Hep G2 (**f**) cells stably expressing Cas9 were transduced with up to 3 different sgRNA targeting ABHD11. Reporter GFP or endogenous HIF-1α levels were measured by flow cytometry (**d**) or immunoblot (**e, f**) respectively after 10-13 days. Endogenous ABHD11 levels were measured by immunoblot and β-actin served as a loading control. (**g**) Reconstitution of mixed KO population of ABHD11 with exogenous ABHD11. HeLa cells expressing Cas9 were transduced with sgRNA targeting ABHD11 as described. Targeted cells were also transduced with exogenous ABHD11 with the PAM site mutated. Cells depleted of PHD2 served as a control for ABHD11 reconstitution. (**h-i**) Quantitative PCR (qPCR) of the HIF-1α target genes (VEGF and CA9) in HeLa cells following ABHD11 depletion by sgRNA (n=3, SEM). sgRNA targeting OGDH and VHL were used as control for HIF-1α activation. (**j**) 5hmC levels are reduced in ABHD11 depleted cells. Genomic DNA was extracted from Hela control or mixed KO populations of ABHD11, LIAS or VHL, and 5hmC levels measured by immunoblot relative to total DNA content (**Figure S1j**). 5hmC levels were quantified using ImageJ. *Ct=control. n=3, Mean ± SEM **p<0.01*.

Both screens identified genes involved in the canonical pathway for HIF stability (VHL, EGLN1 (PHD2)) and 2-OG metabolism (OGDHc components, lipoic acid synthesis pathway), validating the approach (**Figure 1b, c**). Other biological processes that were significantly enriched for sgRNA included intracellular iron metabolism, the mTOR pathway, and transcriptional regulation (**Figure 1b, c**). The reliance of the HIF pathway on these processes is well substantiated and in line with our prior studies using gene-trap mutagenesis in haploid cells^4,15^. In addition to these known pathways, we identified an uncharacterised α/β hydrolase, ABHD11, that was highly enriched for sgRNA in both screens (**Figure 1b, c**).

CRISPR/Cas9 mixed knockout (KO) populations of ABHD11 in HeLa HRE-GFP^ODD^ cells, using three different sgRNA, validated the findings from the screens (**Figure 1d, S1e**). Stabilisation of HIF-1α was observed in multiple cell types (**Figure 1e, 1f, S1f**) and complementation of ABHD11 mixed KO populations with overexpressed ABHD11 restored HIF-1α levels (**Figure 1g**), confirming that ABHD11 loss resulted in HIF-1α accumulation.

We next asked if ABHD11 loss resulted in HIF-1α stabilisation through impaired 2-OG DD activity. PHD function can be readily assessed by measuring HIF-1α prolyl hydroxylation using a HIF prolyl hydroxy-specific antibody. Mixed KO populations of ABHD11 stabilised HIF-1α in a non-hydroxylated form, similar to the HIF-1α stabilisation with DMOG (**Figure S1g**). In contrast, inhibition of the VHL E3 ligase with VH298^20^, which stabilises HIF-1α by preventing ubiquitination and proteasome-mediated degradation showed high levels of hydroxylated HIF-1α (**Figure S1g**). To verify that the decreased prolyl hydroxylation was due to impaired PHD activity, we directly measured prolyl hydroxylation of a recombinant HIF-1α protein in control or ABHD11 deficient lysates^4^. Rapid prolyl hydroxylation was observed with a HeLa control lysate but this was markedly reduced in the ABHD11 depleted cells, similarly to loss of OGDHc function^4^ (**Figure S1h, i**). This PHD inhibition activated a transcriptional HIF response, promoting activation of HIF-1α target genes, VEGF and carbonic anhydrase 9 (CA9), similarly to loss of VHL or OGDH (**Figure 1h, i**).

We also explored whether ABHD11 loss altered the activity of other 2-OG DDs involved in transcription. ABHD11 KO cells showed a marked decrease in total DNA 5-hydroxymethylcytosine (5hmC) levels, similar to those observed when OGDHc function is impaired^4^ (**Figure 1j, S1j**), indicating that Ten-eleven translocation (TET) activity was impaired. However, the activity of selected lysine demethylases (KDMs) were not altered by ABHD11 depletion (**Figure S1k**). Despite these differences between TET and KDM activity, these studies suggested that ABHD11 loss had broader implications for 2-OG DD function, aside from PHDs.

### ABHD11 is required for 2-OG dehydrogenase complex function

Impaired 2-OG DD activity under aerobic conditions suggested that ABHD11 may be involved in 2-OG metabolism. Therefore, we first examined the consequences of ABHD11 loss on 2-OG levels and other TCA cycle intermediates. HeLa cells were depleted of ABHD11 and small molecule metabolites traced by incubating cells with uniformly ^13^C labelled ([U-^13^C_5_]) glutamine, followed by liquid chromatography mass spectrometry (LC-MS) (**Figure 2a**). Cells deficient in OGDH were used as a control to measure perturbations of 2-OG metabolism. ABHD11 depletion resulted in 2-OG accumulation, similarly to OGDH loss (**Figure 2b**). This increase in 2-OG was not due to activation of the HIF response, as we previously demonstrated that PHD2 deficiency does not perturb 2-OG levels^4^. ^13^C tracing confirmed that ABHD11 depletion impaired OGDHc function, as TCA cycle metabolites downstream of the OGDHc were decreased (succinate, fumarate and malate) (**Figure 2c-e**), and cells adapted by showing a shift from oxidative metabolism to reductive carboxylation^4,21^, with a relative decrease in m+4 and m+2 citrate, and an increase in m+5 and m+3 citrate isotopologues (**Figure 2f**).

**Figure 2.**
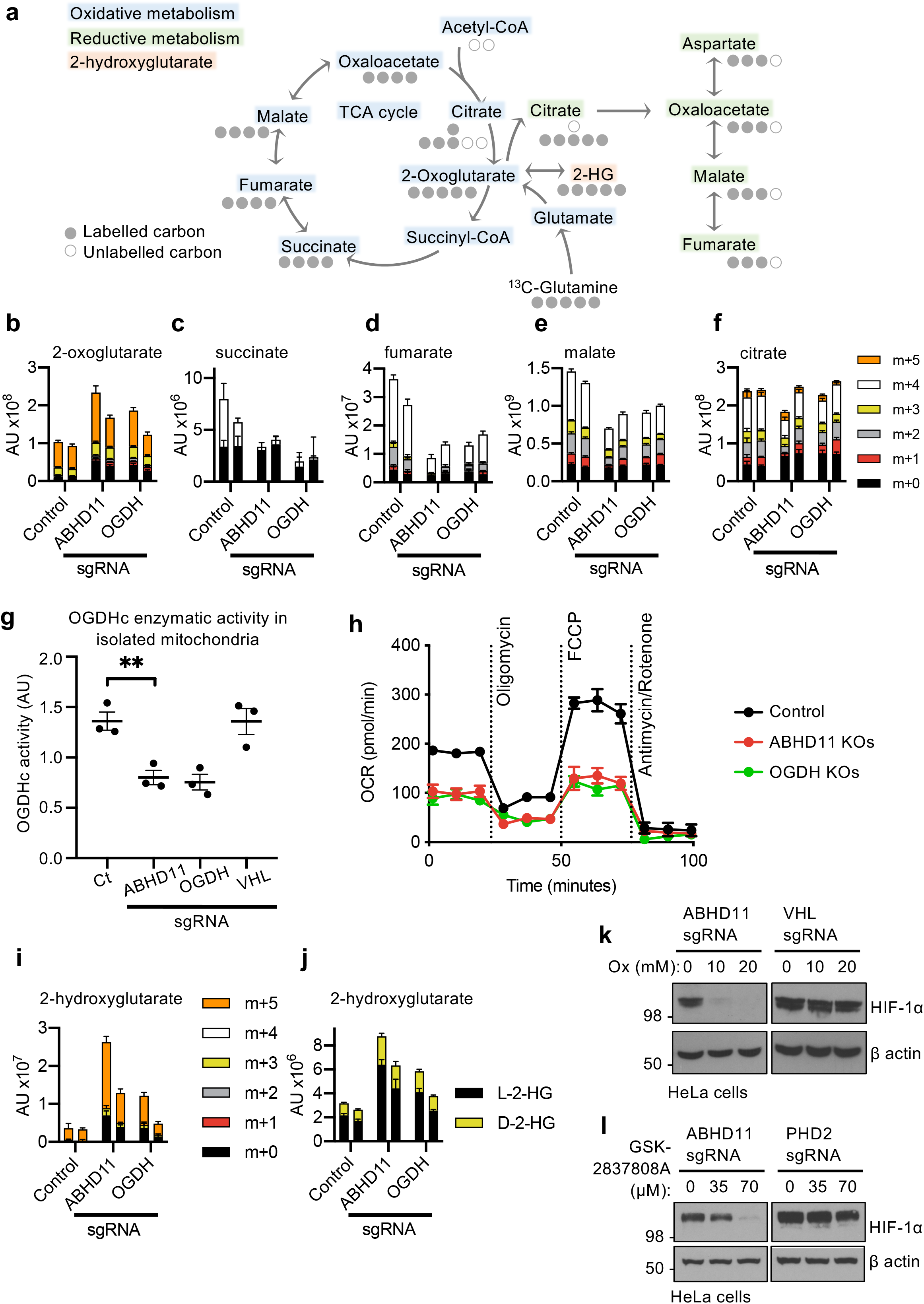
ABHD11 is required for OGDHc function. (**a**) Schematic of the TCA cycle (oxidative metabolism) and reductive carboxylation (reductive metabolism), illustrating the fate of ^13^C carbons upon incubation with [U-^13^C_5_]-glutamine. (**b-g**) Stable isotope tracing of control HeLa cells compared to mixed CRISPR KO populations (sgRNA) of ABHD11 or OGDH incubated with [U-^13^C_5_] glutamine. 2-oxoglutarate (**b**), succinate (**c**), fumarate (**d**), malate (**e**) and citrate (**f**) are shown. The metabolite isotopologues (m+0 to m+5) are indicated. Two biological replicates are shown with five technical repeats. (**g**) OGDHc activity in isolated mitochondria. Mitochondria were extracted from control or mixed CRISPR KO populations of ABHD11, OGDH or VHL HeLa cells and OGDHc activity measured by a redox sensitive colorimetric probe for 2-OG oxidation. Values from three biological experiments are shown. *Mean ± SEM, *p<0.05, **p<0.01*. (**h**) Bioenergetic assays of oxygen consumption rates (OCR) in control, ABHD11 deficient or OGDH deficient HeLa cells (mixed KO populations). ABHD11 and OGDH were depleted as described, and analysed by using a Seahorse XF^e^24 Extracellular Flux Analyzer (4 technical replicates per sample). Three basal measurements were made at 9 minute intervals followed by three measurements per treatment (1μM oligomycin, 1μM FCCP and 1μM antimycin/rotenone). OCR was normalised to total cell number. (**i**) Measurement of 2-hydroxyglutarate (2-HG) levels following [U-^13^C_5_] glutamine stable isotope tracing in control HeLa cells compared to mixed CRISPR KO populations (sgRNA) of ABHD11 or OGDH. Metabolite isotopologues (m+0 to m+5) are indicated. Two biological replicates are shown with five technical repeats. (**j**) Relative quantification of 2-HG enantiomers upon derivatisation with diacetyl-L-tartaric anhydride and LC-MS analysis. (**k, l**) Inhibition of lactate dehydrogenase A (LDHA) in ABHD11 deficient HeLa cells. Mixed CRISPR KO ABHD11, VHL or PHD2 cells were treated with sodium oxamate (Ox) (**k**) or GSK-2837808A (**l**) as indicated for 24 hr. HIF-1α levels were measured by immunoblot.

To substantiate that ABHD11 levels altered OGDHc function, we measured OGDHc enzymatic activity in isolated mitochondria, using a colorimetric assay which detects oxidation of exogenous 2-OG with a redox sensitive probe (**Figure 2g**). OGDHc activity was decreased in ABHD11 deficient mitochondria, similarly to levels observed with depletion of the OGDH subunit (**Figure 2g**). Loss of OGDHc function was not due to HIF stabilisation, as VHL depletion had no effect on OGDHc activity (**Figure 2g**). Bioenergetic profiling also showed that ABHD11 depletion impaired oxygen consumption rates (**Figure 2h**), consistent with a major defect in the TCA cycle and oxidative phosphorylation.

2-OG accumulation can impair 2-OG DD activity through the formation of L-2-HG (**Figure 2a**)^4,6,7,9^. Consistent with this, we observed an accumulation in 2-HG levels following ABHD11 depletion, similarly to OGDH loss (**Figure 2i**). 2-HG predominantly accumulated in its L enantiomeric form, although a small increase in D-2-HG was also observed (**Figure 2j**). Stable isotope tracing with [U-^13^C_5_]glutamine confirmed L-2-HG was derived directly from 2-OG in both the ABHD11 and OGDH deficient cells (m+5 isotopologues)(**Figure 2i**).

Three enzymes are implicated in the formation of L-2-HG from 2-OG: lactate dehydrogenase A (LDHA), malate dehydrogenase 1 (MDH1) and malate dehydrogenase 2 (MDH2)^6,8,9^. However, inhibition of LDHA alone is sufficient to prevent L-2-HG formation^4,7^. Therefore, to confirm that L-2-HG was responsible for decreased 2-OG DD activity, we treated cells with sodium oxamate, which inhibits LDHA as well as decreasing 2-OG formation from glutamine^4,7^, or the selective LDHA inhibitor GSK-2837808A, and measured HIF-1α levels by immunoblot (**Figure 2k, l**). Both treatments restored HIF-1α turnover in ABHD11 deficient HeLa cells. Together, these experiments confirmed that impaired OGDHc function and L-2-HG accumulation was responsible for the decreased PHD activity and activation of the HIF response.

### ABHD11 is a mitochondrial matrix serine hydrolase required for functional lipoylation of the OGDH complex

Conversion of 2-OG to succinyl-CoA by the OGDHc requires decarboxylation and the formation of succinyl intermediate (succinyl-dihydrolipoate), dependent on the cyclical reduction and oxidation of the lipoylated DLST subunit (**Figure 3a**). Therefore, to understand how ABHD11 is required for OGDHc function, we first examined whether protein levels of core OGDHc components or its lipoylation were altered. ABHD11 depletion did not alter total levels of the OGDHc subunits (OGDH, DLST or DLD) in HeLa cells (**Figure 3a, b**). However, using a specific anti-lipoate antibody that detects conjugated lipoamide, we observed a reproducible loss of the faster migrating lipoylated protein species, attributed to the lipoylated DLST subunit of the OGDHc (**Figure 3b, c**). Immunoprecipitation of endogenous DLST confirmed loss of lipoylation following ABHD11 depletion, without altering total DLST levels (**Figure S2a**), and this decreased DLST lipoylation was observed in several cell types (**Figure 3d, 3e**). Furthermore, in contrast to complete disruption of lipoic acid synthesis by LIAS depletion, ABHD11 loss preferentially decreased DLST lipoylation, without altering the other abundantly lipoylated protein within the mitochondria, the DLAT (dihydrolipoamide acetyltransferase) subunit of the pyruvate dehydrogenase complex (PDHc) (**Figure 3b-e**, **S2b**). Indeed, PDHc function, as measured by [U-^13^C_6_] glucose stable isotope tracing, was not impaired in the ABHD11 deficient HeLa cells (**Figure S3a-e**).

**Figure 3.**
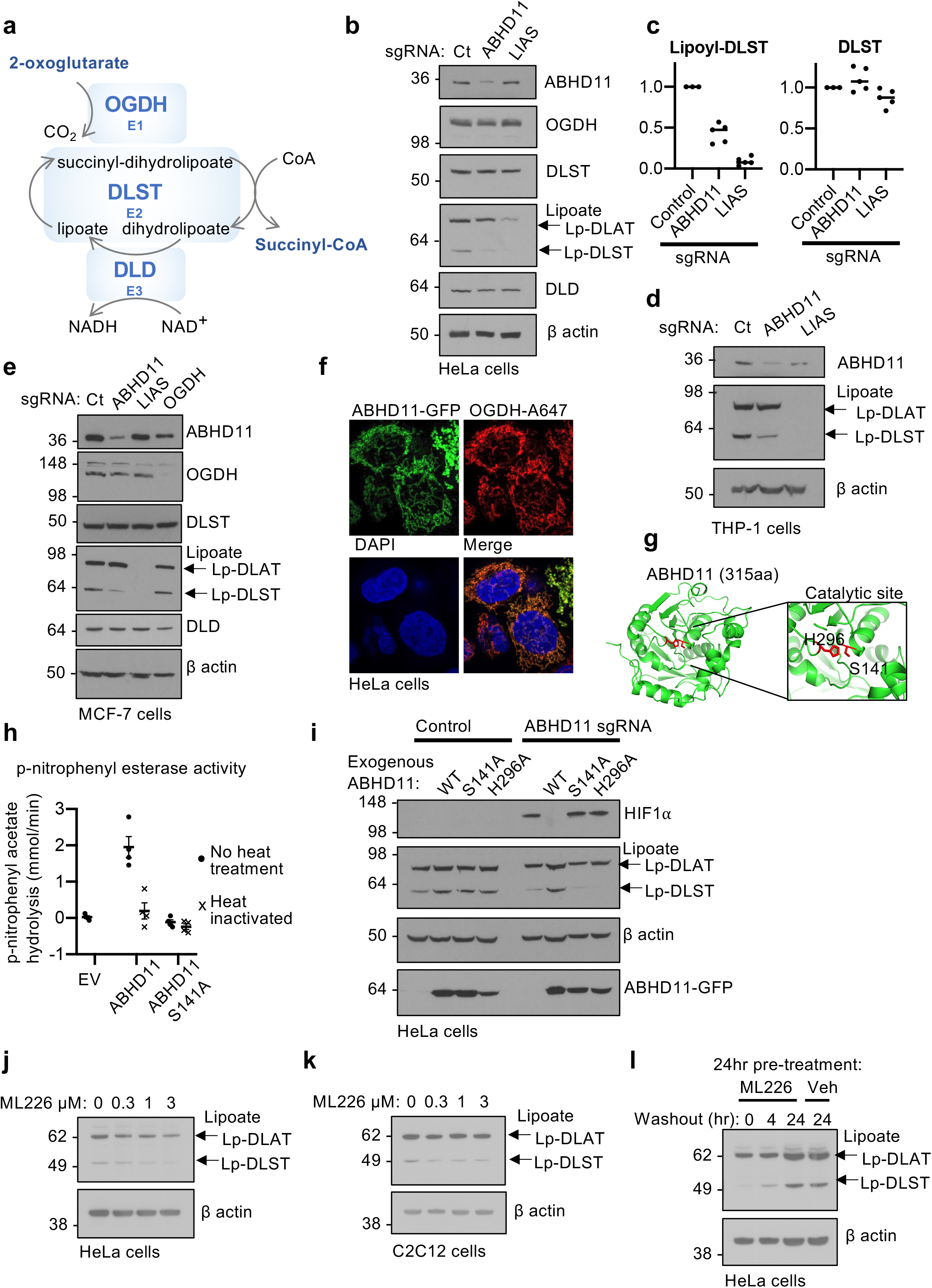
ABHD11 is a mitochondrial matrix serine hydrolase required for functional lipoylation of the OGDHc. (**a**) Schematic of OGDHc function. OGDHc compromises 3 subunits: OGDH (E1) catalyses the oxidation of 2-OG to form the succinyl moiety, releasing carbon dioxide (CO_2_) and reducing the lipoate of DLST to dihydrolipoate, forming a succinyl-dihydrolipoate intermediate through the thiols of lipoate. DLST (E2) then catalyses transfer of the succinyl moiety to Coenzyme A (CoA), forming succinyl CoA and releasing dihydrolipoylated DLST. Dihydrolipoate is oxidised by DLD (E3), forming lipoate, with the free electrons reducing NAD+ to NADH. Thus 2-OG oxidation is coupled to cyclical lipoate reduction/oxidation and NADH formation. (**b, c**) Immunoblot and quantification of OGDHc subunits and lipoylation in ABHD11 or LIAS deficient cells. HeLa cells were transduced with sgRNA targeting ABHD11 or LIAS to generate mixed KO populations, and probed for OGDHc components (**b**). Lipoate (Lp) antibody detects lipoylated proteins. The predominant lipoylated proteins, DLAT and DLST, are indicated. ImageJ quantification of lipoyl-DLST (left) and DLST (right) with five experimental replicates are shown (**c**). *Ct = control, Mean ± SEM.* (**d, e**) Immunoblot OGDHc subunits and lipoylation in ABHD11 or LIAS deficient THP-1 or MCF-7 cells. THP-1 (**d**) or MCF-7 (**e**) cells were transduced with sgRNA targeting ABHD11 or LIAS to generate mixed KO populations, and probed for OGDHc components or lipoate. β-actin served as a loading control. (**f**) Colocalisation of ABHD11 with the mitochondrial matrix protein, OGDH. HeLa cells expressing ABHD11-GFP were fixed in paraformaldehyde. ABHD11 and OGDH subcellular localisation was visualised by immunofluorescence confocal microscopy. *Scale=10μm.* (**g**) *In silico* modelling of ABHD11, indicating the putative catalytic site and key residues S141 and H296 (Phyre2 structural prediction against a template of murine epoxide hydrolase, PDB: 1cr6, and visualised using PyMOL 2.3). (**h**) p-nitrophenyl esterase activity of purified ABHD11-FLAG. Purified wildtype or S141A ABHD11-FLAG were incubated with p-nitrophenyl acetate and hydrolysis measured by rate of increase in absorbance at 405 nm (37°C for 40 min). An empty FLAG vector (EV), that had undergone affinity purification, was used as a control. ABHD11 enzymatic activity was also measured following heat inactivation of the protein (90°C for 5 min). *n=4, Mean ± SEM*. (**i**) Reconstitution of mixed KO population of ABHD11 with exogenous ABHD11-GFP, or enzymatic inactive mutants. (**j, k**) ML266 treatment in HeLa cells (**j**) and C2C12 myoblasts (**k**). Cells were treated for 24 hours with ML226 at the indicated concentrations and lipoylation measured by immunoblot. (**l**) HeLa cells were treated with 1μM ML226, or 0.075% DMSO as a vehicle (veh) control. ML226 was washed out after 24 hr and lipoylation recovery measured by immunoblot.

As our findings suggested that ABHD11 may directly regulate OGDHc lipoylation, we determined if ABHD11 associated with the OGDHc. Endogenous ABHD11 could not be readily detected by immunofluorescence microscopy but exogenously expressed ABHD11 fused to GFP (ABHD11-GFP) localised to the mitochondria (MitoTracker Deep Red) (**Figure S2c, d**) and colocalised with the OGDHc (**Figure 3f**). Endogenous ABHD11 localisation was confirmed using isolated mitochondria and a Proteinase K protection assay. Cytoskeletal and outer membrane space proteins were rapidly lost with the addition of Proteinase K (30 min at 37°C), but ABHD11 levels were unaffected (**Figure S2e**). Furthermore, ABHD11 was retained in the mitoplast fraction, irrespective of proteinase K treatment, consistent with its localisation to the mitochondrial matrix (**Figure S2e**). Endogenous components of the OGDHc (OGDH and DLST) also immunoprecipitated with HA conjugated ABHD11 (ABHD11-HA) (**Figure S2f**), further confirming the OGDHc association.

We next determined if enzymatic activity of ABHD11 altered OGDHc lipoylation. ABHD11 hydrolase activity is predicted to arise from the serine nucleophile motif (GXSXG), but ABHD11 also encodes a putative acyltransferase motif (HXXXXD) found in several other α/β hydrolases with transferase activity^22^ (**Figure 3g**). To first explore whether ABHD11 was an active enzyme, we purified FLAG tagged wildtype and S141A ABHD11 from HEK293T cells (**Figure S4a-d**) and measured hydrolysis of p-nitrophenyl acetate, a validated substrate for generic α/β hydrolase activity ^22,23^. Wildtype ABHD11 hydrolysed the p-nitrophenyl ester, but this was lost with the S141A mutant or following heat treatment (**Figure 3h**), confirming that ABHD11 was a functional serine hydrolase.

Complementation studies were used to determine whether the enzymatic activity of ABHD11 was required for lipoylation of the OGDHc. Exogenous wildtype or S141A mutant ABHD11 was expressed in mixed ABHD11 KO populations and lipoylation levels measured by immunoblot. DLST lipoylation was restored with the wildtype ABHD11 but not with the S141A mutant (**Figure 3i**). Furthermore, HIF-1α levels were only reduced to basal levels by reconstituting with wildtype ABHD11 but not the nucleophile mutant.

In addition to a serine hydrolase domain, ABHD11 encodes a putative acyltransferase motif (HXXXXD)^22^. A histidine to alanine mutation in this domain (ABHD11 H296A) failed to reconstitute OGDHc lipoylation in mixed ABHD11 null cells (**Figure 3i**), similarly to the S141A mutation, but we were unable purify this mutant to test the enzyme activity of this mutant *in vitro*. However, both ABHD11 S141A and H296A localised to the mitochondria (**Figure S2c, d**), confirming that their failure to restore OGDHc lipoylation was not due to a trafficking defect.

Lastly, to confirm the enzymatic requirement of ABHD11 in OGDHc lipoylation, we used a highly selective covalent inhibitor of ABHD11, ML226, that was initially developed as a tool for screening serine hydrolases^24^. ML226 treatment inhibited ABHD11 p-nitrophenyl ester hydrolysis *in vitro* (**Figure S2g**) and decreased OGDHc lipoylation in HeLa cells (**Figure 3j**) and cultured myoblasts (C2C12) (**Figure 3k**). In addition, DLST lipoylation recovered after ML226 washout in cells, indicating OGDHc lipoylation can be restored (**Figure 3l**). Thus, a functional OGDHc requires the hydrolase activity of ABHD11.

### ABHD11 prevents the formation of lipoyl adducts by lipid peroxidation products

The finding that ABHD11 loss showed a selective loss of DLST lipoylation was unexpected, as prior genetic studies of lipoate conjugation had not shown a requirement for an additional enzyme^25^. Furthermore, we confirmed that ABHD11 loss differed to depletion of other components of the lipoic acid synthesis pathway by generating CRISPR/Cas9 mixed KO populations of the key enzymes involved (**Figure S5a, b**). Lipoyl(octanoyl) transferase 2 (LIPT2), LIAS, and lipoyltransferase 1 (LIPT1) all reduced DLAT and DLST lipoylation in HeLa cells to a similar level, but only ABHD11 showed a selective loss of DLST lipoylation (**Figure S5b**). This preferential decrease in DLST lipoylation following ABHD11 loss argued against a general role for ABHD11 in lipoyl synthesis, and while it remained possible that ABHD11 was required for the final catalysis of DLST lipoylation, prior genetic studies suggested that LIPT1 was sufficient for this step^25–27^.

Rather than acting as a conjugating enzyme, we hypothesized that ABHD11 may instead be involved in maintaining a functional lipoate moiety on the OGDHc complex. Therefore, we used mass spectrometry to directly determine how ABHD11 activity altered DLST lipoylation. Immunoprecipitated DLST was treated with a reducing agent and then incubated with N-ethylmaleimide (NEM), forming an NEM-lipoyl conjugate, which had previously been shown to aid detection of the lipoate moiety^28^ (**Figure 4a, S5c**). Interestingly, NEM treatment prevented detection of immunoprecipitated lipoylated DLST by immunoblot (**Figure S5c**), demonstrating that the anti-lipoate antibody only detected the functional lipoate moiety and not the NEM-modified form, consistent with our hypothesis that the apparent loss of DLST lipoylation in ABHD11 deficient cells may due to the formation of lipoyl adducts. We next measured levels of DLST lipoylation (NEM-lipoyl) by label-free quantification on immunoprecipitated DLST from wildtype HeLa cells or those deficient in LIAS or ABHD11 (**Figure 4a, b, S5d**). To account for potential differences in DLST protein abundance around the lipoylated region (DK*TSVQVPSPA), we normalised these peptides to the sum of all DLST peptide label-free quantification values. Approximately 50% of the DK*TSVQVPSPA DLST peptide in wildtype cells was modified with lipoate compared to the unmodified form and as expected, nearly all the lipoate detected was modified with NEM (**Figure 4b**). DLST lipoylation was nearly completely lost in the LIAS deficient cells, with the majority of the DLST lipoylated peptide region found to be unmodified (**Figure 4b**), confirming that this approach could readily identify a defect in lipoyl synthesis and conjugation. However, ABHD11 deficiency did not result in an accumulation of the unmodified DK*TSVQVPSPA peptide, which could have been expected with a defect in conjugation. Instead, both the unconjugated or NEM-lipoyl DK*TSVQVPSPA peptide were barely detectable, with a 10-fold decrease in abundance compared to the control or LIAS null cells (**Figure 4b**). This decrease was not due to less total DLST, as DK*TSVQVPSPA levels were normalised to other DLST peptides upstream or downstream of the lipoylated region. Therefore, a modification of the DK*TSVQVPSPA peptide of undefined mass accounted for the apparent decrease in peptide abundance. Common post-translational modifications (e.g. ubiquitination, phosphorylation or acetylation), combinations of modifications, or known DLST intermediates (e.g. succinyl-dihydrolipoamide, acyl-dihydrolipoamide or S-glutathionylation) (**Supplementary File S2**) did not account for the peptide loss of the lipoylated DLST region, suggesting the formation of lipoyl adducts that were not detectable by mass spectrometry.

**Figure 4.**
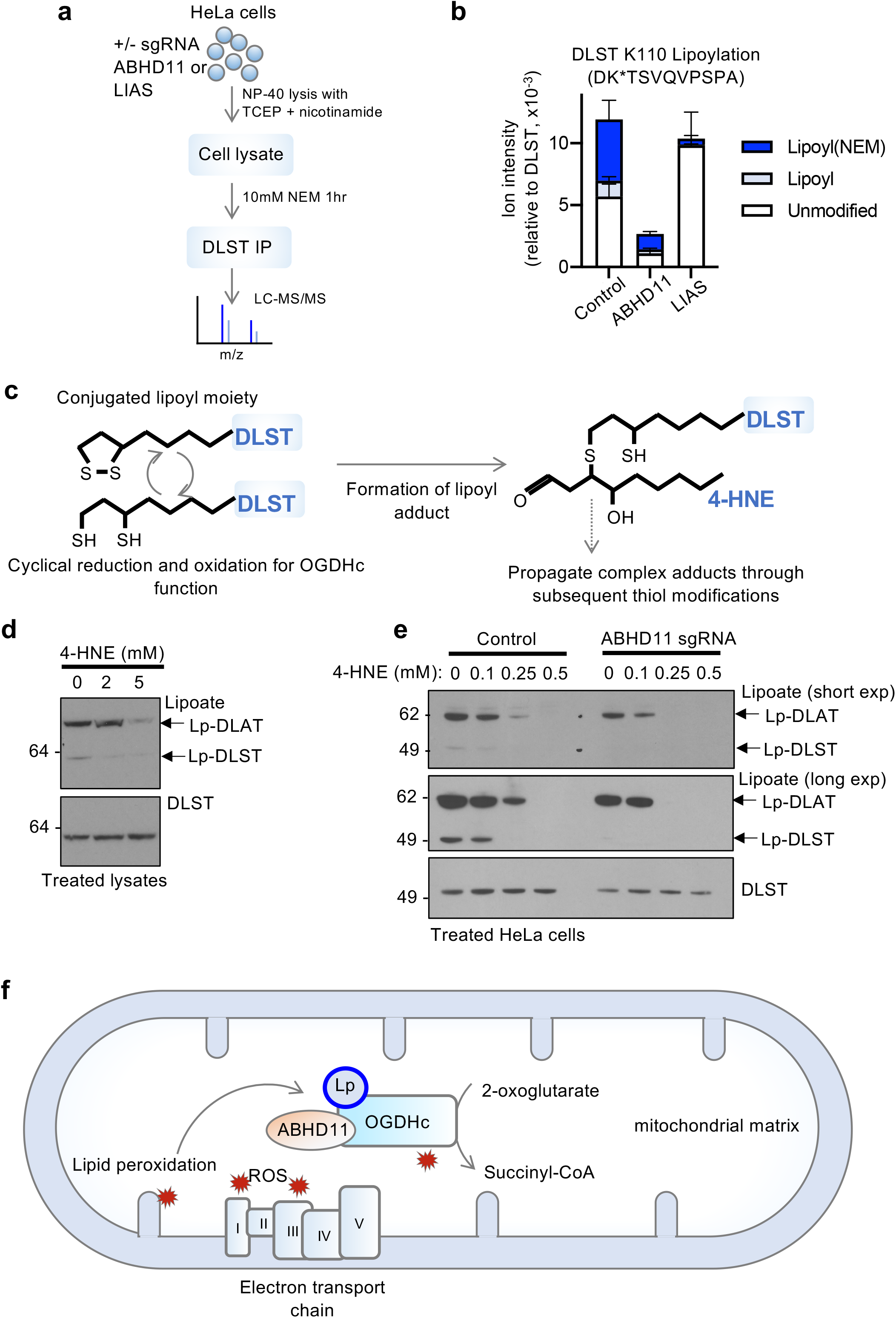
ABHD11 prevents the formation of lipoyl adducts by lipid peroxidation products. (**a, b**) Mass spectrometry analysis of DLST lipoylation. DLST was immunoprecipitated from HeLa control, ABHD11 deficient or LIAS deficient cells, and treated with NEM to modify the free thiols and maintain the lipoyl moiety in a reduced state. After SDS-PAGE, protein samples were digested with Asp-N protease and analysed by LC-MS/MS. (**b**) Normalised level of lipoylated DLST peptide compared to DLST reference peptide. Relative levels of the unmodified, NEM-dihydrolipoamide, and lipoamide DLST peptide are shown. *n=3, Mean ± SEM.* (**c**) Schematic of normal cyclical reduction and oxidation of the lipoyl moiety on DLST (left). The reduced dihydrolipoamide can react with lipid peroxidation products (e.g. 4-HNE) to form lipoyl adducts through the free thiols, which may also propagate (right). (**d, e**) Effect of 4-HNE treatment on lipoylation. HeLa cells lysates were treated with 4-HNE at the indicated concentrations for 60 minutes (50°C) and immunoblotted for lipoylated proteins or total DLST (**d**). Control or ABHD11 deficient HeLa cells were treated with 4-HNE for 90 minutes (37°C), lysed and immunoblotted for lipoylation and total DLST (**e**). Short and long lipoate antibody photographic exposures are shown. (**f**) Working model for functional lipoylation of the OGDHc. Cyclical reduction and oxidation of lipoate is required for OGDHc catalytic activity, but this process can be disrupted by lipid peroxidation products (e.g. 4-HNE) formed by ROS generated by the OGDHc or by the electron transport chain. ABHD11 maintains DLST lipoylation, reducing the levels of lipoyl adducts, facilitating TCA cycle function and 2-OG dependent signalling.

The lipoamide moiety is highly sensitive to attack by lipid peroxidation products, which disrupt OGDHc catalysis by preventing the cyclical oxidation and reduction of the lipoyl conjugate^29,30^ (**Figure 4c**). While the exact nature of the lipoyl adduct formed in ABHD11 deficient cells was unclear, we examined whether exogenous 4-hydroxy-2-nonenal (4-HNE), a common lipid peroxidation product known to interfere with OGDHc function^29–31^, altered DLST lipoylation, similarly to ABHD11 depletion. 4-HNE treatment of cell lysates preferentially decreased detection of DLST lipoylation by immunoblot (**Figure 4d**), consistent with the formation of lipoyl adducts preventing binding to the antibody. DLAT lipoylation was only affected at high concentrations (5 mM) of 4-HNE (**Figure 4d**), suggesting that DLAT may be more resistant to lipoyl adduct formation than DLST, consistent with our findings that ABHD11 loss preferentially effects the OGDHc.

Finally, to substantiate that ABHD11 protected against to the formation of these lipoyl adducts, we measured whether ABHD11 loss or inhibition made the OGDHc more susceptible to lipid peroxidation damage. Control or ABHD11 depleted HeLa cells were treated with low concentrations of 4-HNE and lipoylation detected by immunoblot. 4-HNE decreased DLST and DLAT functional lipoylation in a concentration dependent manner, with loss of DLST lipoylation prior to DLAT (**Figure 4e**), as observed in the cell extracts (**Figure 4d**). However, ABHD11 deficient cells were more susceptible to 4-HNE treatment compared to the control cells, with complete loss of all functional lipoylation was observed with 0.25 mM 4-HNE and selective loss of DLST lipoylation at 0.1 mM (**Figure 4e**). Similar findings were observed with the specific mitochondrial ROS inducing agent, mitoparaquat (MitoPQ)^32^, and overexpression of ABHD11 inactive mutants, ABHD11 S141A-GFP or ABHD11 H296A-GFP, which competed with endogenous ABHD11 to also show an increase lipoyl-adduct formation following 4-HNE treatment (**Figure S5e**). Thus, ABHD11 maintains OGDHc function by preventing lipoyl adduct formation in the context of increased ROS and lipid peroxidation products (**Figure 4f**).

## Discussion

This study identifies ABHD11 as a mitochondrial enzyme required for OGDHc function, and to our knowledge, is the first example of a mitochondrial specific response that maintains TCA cycle integrity by counteracting the damage caused by lipid peroxidation products (**Figure 4f**). Moreover, we demonstrate that ABHD11 inhibition allows 2-OG metabolism to be modulated in multiple cells and in a reversible manner, with potential broad implications for altering cell fate-decisions and manipulating 2-OG abundance in tumours.

The selective loss of lipoylated DLST following ABHD11 depletion initially suggested that it may be necessary for OGDHc lipoate conjugation. However, a requirement for ABHD11 in lipoate synthesis had not been previously observed ^13,25,33^, and LIPT1 deletion or human loss of function mutations prevent PDHc and OGDHc lipoylation^25–27^. Instead, we found an absence of both the modified and unmodified lipoylated region of DLST by mass spectrometry (**Figure 4b, c**), consistent with the formation of a lipoyl adduct.

Lipid peroxidation products (hydroxyalkenals, such as 4-HNE), arise from free radical propagation through phospholipids^30^, and can easily react with thiol groups, inactivating 2-oxoacid dehydrogenases by forming lipoyl adducts^29^. In addition, the hydrophobicity and complex nature of these adducts^31^ precludes their easy detection by mass spectrometry, which would account for the apparent loss of the DLST DK*TSVQVPSPA peptide that we observed (**Figure 4b**) and failure to detect 4-HNE modified forms. However, consistent with prior reports, we find that the OGDHc is more susceptible to 4-HNE adduct formation than the lipoylated PDHc proteins^34^, which may account for the apparent specificity of ABHD11 for DLST.

Lipoic acid has been traditionally described as an essential cofactor for 2-oxoacid dehydrogenases, but only 50% of DLST in HeLa cells was observed to be lipoylated in resting cells (**Figure 4b**), and OGDHc lipoylation was rapidly stored after washout of ML226, suggesting a reserve capacity to alter lipoate levels and increase OGDHc activity (**Figure 3j, k**). These findings show that lipoylation is a dynamic modification that must be maintained, which is further supported by recent observations that SIRT4 act as a lipoamidase, altering PDHc function^28^, and that increased lipoylation can enhance brown adipose tissue function, decreasing age-associated obesity^35^. Therefore, modulating lipoylation through ABHD11 activity provides a novel approach to manipulating 2-OG metabolism. Moreover, these studies extend the role of lipoylation beyond an enzymatic cofactor, to a dynamic modification that couples the mitochondrial redox environment to a transcriptional adaptive response mediated by 2-OG and oxygen sensitive enzymes.

## Methods

### Cell lines and reagents

HeLa, MCF-7, C2C12 and HEK293T cells were maintained in DMEM (Sigma D6429), and THP-1 cells maintained in RPMI-1640 (Sigma R8758), both supplemented with 10% foetal calf serum (Sigma P4333) and 100 units/ml penicillin with 100 μg/ml streptomycin, in a 5% CO_2_ incubator at 37°C. HeLa HRE-GFP^ODD^ cells were produced as described previously ^4^. All cell lines were authenticated (Eurofins). Hypoxia treatments were performed at 1% oxygen, 5% CO_2_ in a H35 Hypoxystation (Don Whitley Scientific).

Full details of reagents and antibodies used are shown in **Table S1.**

### Constructs

#### Lentiviral CRISPR sgRNA

CRISPR sgRNAs were cloned into a lentiviral sgRNA expression vector pKLV-U6gRNA(BbsI)-PGKpuro2ABFP ^36^. All sgRNA sequences are detailed in **Table S1.**

#### Expression plasmids

ABHD11 constructs were generated from the I.M.A.G.E. cDNA clone (IRATp970F0688D, Source Bioscience), cloned into the pHRSIN pSFFV backbone with pGK-blasticidin resistance (a gift from Paul Lehner), using NEBuilder HiFi (NEB). Prior to assembly, silent mutations were introduced inside the sequence targeted by ABHD11 sgRNA 2, using PCR primers detailed in **Table S2**. Mutations of catalytic residues serine 141 to alanine and histidine 296 to alanine of ABHD11 was created using NEBuilder HiFi, with primers detailed in **Table S2**. Lentiviral expression vectors (pHRSIN) for ABHD11, S141A ABHD11 and H296A ABHD11 with C-terminal eGFP tags or HA tags were created using NEBuilder HiFi (NEB). ABHD11 was also cloned into a transfection vector, pCEFL 3xFLAG mCherry vector, encoding a C-terminal 3X FLAG tag and mCherry under a separate promoter (a gift from David Ron)^37^, using Gibson Assembly (NEB) and NEBuilder HiFi.

### CRISPR/Cas9 sgRNA pooled libraries

The Human Brunello library was a gift from David Root and John Doench^18^ (Addgene #73178). The Toronto human knockout pooled library (TKO) Version 1 (Addgene #1000000069) was a gift from Jason Moffat^19^.

### Preparation of lentivirus and transductions

HEK293T cells were transfected using TransIT-293 (Mirus) according to the manufacturer’s protocol. For small scale experiments, lentivirus was produced in 6-well plates containing 2 ml media, using 2 μg DNA (DNA was mixed in a 3:2:4 ratio of the relevant expression plasmid, pCMV-dR8.91 (gag/pol) and pMD.G (VSVG)^38^. Viral supernatant was harvested after 48 hours, passed through a 0.45 μm filter, and frozen at −80°C.

Cells were transduced with lentivirus by adding an appropriate volume of thawed viral supernatant. In the case of single-gene knockdown in HeLa cells, 250 μl of virus with 5×10^4^ cells in a 24-well plate made up to 1 ml media. For the screens, a titration of increasing volumes of virus was used, with 10^6^ HeLa cells in a 6-well plate. Cell plates were centrifuged for 1 hour at 37°C at 750 *g* immediately after addition of virus.

### Whole genome CRISPR/Cas9 forward genetic screens

HeLa HRE-GFP^ODD^ cells were transduced with Streptococcus pyogenes Cas9 (pHRSIN-FLAG-NLS-CAS9-NLS-pGK-Hygro)^39^ and selected for Cas9 expression using hygromycin. 5×10^7^ (Brunello) or 10^8^ (TKO) HeLa HRE-GFP^ODD^ cells were transduced with the appropriate volume of pooled sgRNA virus (multiplicity of infection (MOI) of approximately 0.3), maintaining at least 150-fold sgRNA coverage. After 30 hours, cells were treated with puromycin 1μg/ml for 5 days. Representation was maintained throughout the screen such that no selection event occurred where the library was cultured at fewer than 200 times the number of sgRNA sequences in the library. The library was pooled immediately before any selection event.

Fluorescence-activated cell sorting (FACS) was performed by harvesting 10^8^ cells, washing the cells in PBS, and then resuspending them in PBS containing 2% fetal calf serum and 10 mM HEPES (Sigma H0887). Cells were sorted using an Influx cell sorter (BD); GFP-high cells were chosen in a gate set at one log_10_ unit above the mode of the untreated population.

Genomic DNA was extracted using a Gentra Puregene Core kit (Qiagen). Lentiviral sgRNA inserts were amplified in a two-step PCR (with Illumina adapters added on the second PCR), as previously described^39^. For the TKO screen, the forward inner PCR and sequencing primers were modified (**Table S2**).

Sequencing analysis was performed by first extracting the raw sequencing reads, trimming the first 20 bp, and aligning against the appropriate sgRNA library using Bowtie^40^. Read counts for each sgRNA were compared between conditions, and Benjamini-Hochberg false discovery rates for each gene calculated, using MAGeCK^41^ (**Supplementary File S1**). The analysis presented compares DNA extracted following the second sort to an unsorted DNA library taken at the same timepoint.

### Immunoblotting and immunoprecipitation

Cells were lysed in an SDS lysis buffer containing 2% SDS, 50 mM Tris pH 7.4, 150 mM NaCl, 1 mM dithiothreitol, 10% glycerol and 1:200 Benzonase nuclease (Sigma), for 15 minutes at room temperature, then heated at 90°C for 5 minutes. Proteins were separated with SDS-PAGE electrophoresis, transferred to a PVDF membrane, and probed using appropriate primary antibodies and a secondary with HRP conjugate. Densitometry measurements were made using ImageJ^42^.

To identify protein interactions with ABHD11, HeLa cells lentivirally transduced with ABHD11 with a C-terminal HA tag were lysed in a buffer containing 100 mM Tris pH 8.0, 140 mM NaCl, 1% IGEPAL CA-630 (Sigma), 1 mM PMSF (Sigma P7626) and cOmplete Protease Inhibitor Cocktail (Roche). After centrifugation at 17,000 *g*, the supernatant was pre-cleared using Sepharose CL-4B (GE Heathcare) and then incubated with EZView HA Red Anti-HA Affinity Gel (Sigma E6779) overnight on a rotator. Resins were washed with Tris-buffered saline containing 0.1% IGEPAL CA-630, and a further 2 washes with Tris-buffered saline. Proteins were eluted using an SDS lysis buffer (4% SDS, 100 mM Tris pH 7.4, 300 mM NaCl, 2 mM dithiothreitol, and 20% glycerol) heated at 90°C for 5 minutes, and separated using SDS-PAGE.

### Confocal microscopy

Mitochondrial labelling was performed using MitoTracker Deep Red FM (Thermo M22426). Cells were cultured overnight on a 1 cm glass coverslip, incubated with 250 nM Mitotracker Deep Red FM for 40 minutes, and after washing with PBS fixed with 4% paraformaldehyde for 20 minutes. Cells were mounted to slides using ProLong Gold Antifade Mountant with DAPI (Thermo).

For immunofluorescence microscopy of OGDH, cells were cultured overnight on a 1 cm glass coverslip. After washing with PBS, cells were fixed with 4% paraformaldehyde, permeablised (0.3% Triton X-100, 3% bovine serum albumin) and stained for OGDH (primary antibody at 1:100 dilution for 30 minutes, Alexa Fluor secondary at 1:1000 (Thermo A21245)). Cells were mounted to slides using ProLong Gold Antifade Mountant with DAPI. Slides were imaged using an LSM880 confocal microscope (Zeiss).

### Quantitative PCR

Quantitative PCR was performed as described previously^4^. Briefly, total RNA was extracted using the RNeasy Plus minikit (Qiagen) and reverse transcribed using Super RT reverse transcriptase (HT Biotechnology Ltd). PCR was performed on the ABI 7900HT Real-Time PCR system (Applied Biosystems) using SYBR Green Master mix (Applied Biosystems). Reactions were performed with 125ng of template cDNA. Transcript levels of genes were normalised to a reference index of housekeeping genes (GAPDH and RPS2).

### Measurements of 2-OG dependent dioxygenase activity

#### In vitro HIF1α prolyl hydroxylation assay

The hydroxylation activity of HeLa cell lysates against a His-tagged protein corresponding to residues 530-652 of human HIF-1⍺ protein was performed as previously described^4,15^. Images were quantified using ImageJ, and are presented as the ratio of densitometry of hydroxylated to total HIF-1⍺^ODD^ peptide at the 15 minute timepoint, normalised to PHD2.

#### 5hmC dot blot assay

Genomic DNA was extracted from HeLa cells using a Gentra Puregene kit (Qiagen), and dot blotting for 5hmC levels performed as previously described ^4^.

#### KDM panel

Cells were lysed in SDS lysis buffer and probed for selected H3 methylation marks.

### Mitochondrial protease protection assay

Mitochondria from 10^7^ HeLa cells were extracted using a Qproteome Mitochondria Isolation Kit (Qiagen). After the final wash with mitochondria storage buffer, mitochondria were divided into tubes, pelleted by centrifugation at 6,000 *g* and resuspended in either 10 mM Tris-HCl pH 8.0 with 250 mM sucrose (for whole mitochondria), or 10 mM Tris-HCl pH 8.0 (for mitoplasts), to a final protein concentration of 1 mg/ml. Proteinase K (Sigma P2308) was added to a final concentration of 12 or 24 μg/ml, based on methods previously described^43^. Following incubation at 37°C for 30 minutes, the reaction was quenched with 1mM PMSF. Mitochondria or mitoplasts were then pelleted again by centrifugation at 6,000 *g*, lysed in SDS buffer and analysed by immunoblot.

### Purification of ABHD11 from HEK293T cells

HEK293T cells were transfected with the pCEFL-ABHD11 −3XFLAG tag plasmid. Briefly, cells were seeded in a 14 cm dish at 70% confluency and transfected using 270 μg polyethylenimine (Sigma) with 22.5 μg DNA in 6 ml Opti-MEM. Cells were harvested after 48 hours, and lysed in TBS buffer (100 mM Tris-HCl pH 8.0, 140 mM NaCl) with 1% Triton X-100 and cOmplete Protease Inhibitor Cocktail (Roche). After centrifugation at 17,000 *g*, the supernatant was pre-cleared using Sepharose CL-4B (GE Heathcare) and incubated overnight with FLAG M2 antibody conjugated beads (Sigma). Following 5 washes with TBS, ABHD11-FLAG was eluted using 100 mg/l 3xFLAG peptide (Sigma F4799), filtered using a 0.22 μm PVDF filter and separated using a Superdex 75 10/300 column on an Äkta-Pure liquid chromatography system. 500 μl fractions were collected and protein content visualised by SDS-PAGE and Coomassie staining. Protein identity was confirmed by LC-MS/MS.

### Liquid chromatography mass spectrometry

Samples were reduced, alkylated and digested in-gel using either trypsin, GluC or AspN. The resulting peptides were analysed by LC-MS/MS using an Orbitrap Fusion Lumos coupled to an Ultimate 3000 RSLC nano UHPLC equipped with a 100 μm ID × 2 cm Acclaim PepMap Precolumn (Thermo Fisher Scientific) and a 75 μm ID × 50 cm, 2 μm particle Acclaim PepMap RSLC analytical column. Loading solvent was 0.1% formic acid with analytical solvents A: 0.1% formic acid and B: 80% acetonitrile + 0.1% formic acid. Samples were loaded at 5 μl/minute loading solvent for 5 minutes before beginning the analytical gradient. The analytical gradient was 3-40% B over 42 minutes rising to 95% B by 45 minutes followed by a 4 min wash at 95% B and equilibration at 3% solvent B for 10 minutes. Columns were held at 40°C. Data was acquired in a DDA fashion with the following settings: MS1: 375-1500 Th, 120,000 resolution, 4×10^5^ AGC target, 50 ms maximum injection time. MS2: Quadrupole isolation at an isolation width of m/z 1.6, HCD fragmentation (NCE 30) with fragment ions scanning in the Orbitrap from m/z 110, 5×10^4^ AGC target, 100 ms maximum injection time. Dynamic exclusion was set to +/- 10 ppm for 60 s. MS2 fragmentation was trigged on precursors 5×10^4^ counts and above.

Raw files were processed using PEAKS Studio (version 8.0, Bioinformatics Solutions Inc.). Searches were performed with either trypsin, GluC or AspN against a Homo sapiens database (UniProt reference proteome downloaded 26/01/18 containing 25,813 sequences) and an additional contaminant database (containing 246 common contaminants). Variable modifications at PEAKS DB stage included oxidation (M) and carbamidomethylatation with 479 built in modifications included at PEAKS PTM stage.

### p-Nitrophenyl ester hydrolysis assay

Hydrolase activity of ABHD11-FLAG (or a heat-inactivated control made by incubation at 90°C for 5 minutes) was assayed by incubation in an assay buffer containing 50 mM Tris-HCl pH 7.4, 150 mM NaCl, 0.01% bovine serum albumin, 1.4% methanol, and 500 μM p-nitrophenyl acetate (Sigma N8130). 1.5 μg enzyme was added to 200 μl assay buffer, and the formation of p-nitrophenol assayed using a Clariostar plate reader (BMG Labtech), recording absorbance at 405 nm while incubating at 37°C for 40 min. The rate of formation of p-nitrophenol was calculated from the slope of the absorbance curve, subtracting the slope of a blank containing only assay buffer and substrate, and calibrated against a standard curve of p-nitrophenol.

### OGDHc activity assay

OGDHc activity was measured in whole cell lysates using a Biovision ketoglutarate dehydrogenase activity assay (Biovision K678), according to the manufacturer’s protocol. HeLa cells were lysed by three freeze-thaw cycles followed by passing 10 times through a 26-gauge needle. Activity was subtracted from a background control containing cell lysate but no substrate.

### Mitochondrial bioenergetics assay

Dynamic measurements of oxygen consumption rate and extracellular acidification were recorded using a SeaHorse XFe24 (Agilent). HeLa cells were seeded 24 hours beforehand at 1.5 × 10^4^ cells/well, and assayed using the manufacturer’s Mito Stress Test protocol.

### Stable isotope tracing by Liquid Chromatography Mass Spectrometry (LC-MS)

Samples were analysed as described previously^4^. Briefly, HeLa cells were seeded in 5 replicates in 6-well plates 27 hours prior to metabolite extraction, with a sixth well per condition used for cell count. 24 hours prior to extraction, media was changed to either DMEM without L-glutamine (Sigma D6546) supplemented with 10% FCS and 4 mM [U-^13^C_5_]L-glutamine (Cambridge Isotopes CLM-1822), or DMEM without glucose (Gibco 11966-025) supplemented with 10% FCS, 1 mM sodium pyruvate and 25 mM [U-^13^C_6_]D-glucose (Cambridge Isotopes CLM-1396). Metabolites were extracted on dry ice, after washing with ice cold PBS, with 1ml per 10^6^ cells of extraction buffer, containing 50% methanol, 30% acetonitrile, 20% water and 100 ng/ml HEPES. To quantify the two enantiomers of hydroxyglutarate, a subset was derivatised using 50 mg/ml diacetyl-L-tartaric anhydride in 20% acetic acid/80% acetonitrile, as described previously ^4^.

### Immunoprecipitation and mass spectrometry detection of lipoylation

HeLa cells were lysed on ice for 30 min in a HEPES buffer (150 mM NaCl, 50 mM HEPES pH 7.0, 1% IGEPAL CA-630, 1 mM nicotinamide (Sigma), 5 mM tris(2-carboxyethyl)phosphine hydrochloride (Sigma), 1X cOmplete Protease Inhibitor Cocktail (Roche), and 1 mM PMSF). Lysates were centrifuged at 16,900 *g* for 10 min. For LC-MS experiments, free thiols were blocked by addition of 10 mM N-ethyl maleimide (Sigma) and incubated on a rotator at 4°C for 1 hour.

Lysates were immunoprecipitated by incubation with protein G beads (GE 17-0168-01), firstly for 1 hour to pre-clear, and then overnight with the DLST or DLAT antibodies (**Table S1**). Resins were washed four times with Tris-buffered saline containing 0.1% IGEPAL CA-630, followed by three washes with Tris-buffered saline. Protein was eluted using 2% SDS, 50 mM Tris pH 7.4, 150 mM NaCl, 1 mM dithiothreitol and 10% glycerol, and incubated at 90°C for 5 minutes. The proteins were then separated using SDS PAGE (Thermo NP0335), and visualised using SimplyBlue SafeStain (Invitrogen).

For mass spectrometry analysis, in-gel AspN digest and sample analysis were performed as previously described. To identify possible modifications of DLST K110, raw files were processed using PEAKS Studio (version 8.0, Bioinformatics Solutions Inc.) with the following parameters: AspN digestion; Human database (UniProt reference proteome downloaded 18 Dec 2018 containing 21066 proteins) with additional contaminant database (containing 246 common contaminants); oxidation and carbamidomethylation as variable modifications at the PEAKS DB stage. The data were processed twice, with different variable modifications searched at the PEAKS PTM stage, either with 485 PEAKS built-in modifications listed in **Supplementary File S2**(sheet A) and 34 custom modifications listed in **Supplementary File S2**(sheet B).

XIC were obtained from Thermo Xcablibur Qual Browser (4.0.27.13). Mass ranges were limited to 564.79-564.80 m/z, 658.81-658.82 m/z and 784.87-784.88 m/z for the unmodified, lipoylated and 2x NEM lipoylated peptides respectively.

Label-free quantitation values were obtained by processing raw files with MaxQuant (version 1.6.6.0) with the following parameters: specific AspN digestion; Human database (UniProt reference proteome downloaded 18 Dec 2018 containing 21066 proteins); oxidation, lipoylation, 2x NEM lipoylation, N terminal acetylation as variable modifications; carbamidomethylation as a fixed modification; label-free quantification enabled. Label-free quantification values were normalised to the sum of DLST peptides label-free quantification values.

### ABHD11 structural prediction

A structural model of ABHD11 was obtained using the NCBI reference sequence for ABHD11 transcript variant 1 (NP_683710.1), modelled with Phyre2 against a template of murine epoxide hydrolase (PDB: 1cr6) ^44^ and visualised using PyMOL 2.3 (Schrödinger, LLC). The mitochondrial targeting sequence was mapped with the MitoFates prediction tool^45^.

### Quantification and Statistical Analysis

Statistical analysis of the screens was performed using MAGeCK version 0.5.5^41^, testing the sgRNA read counts obtained following the second sort against sgRNA read counts obtained from unsorted cells lysed at the same timepoint. Quantification and data analysis of other experiments are expressed as mean ± SEM and P values were calculated using two-tailed Student’s t-test for pairwise comparisons, unless otherwise stated. Metabolomic samples were blinded and randomised prior to their evaluation.

### Data and Software Availability

SgRNA read count tables from CRISPR/Cas9 genetic screens are shown in **Supplementary File S1**. Proteomics data has been deposited in the PRIDE archive (https://www.ebi.ac.uk/pride/archive/) (submission in process) and the results and analysis are also available in **Supplementary File S2**.

## Author Contributions

Conceptualization, PSJB and JAN; Methodology, PSJB, BMO, ASHC, JWH, RA, CF and JAN; Investigation, PSJB, BMO, ASHC, JWH, RA, and JAN; Writing – original draft, PSJB and JAN; Writing – reviewing and editing, PSJB, ASHC, CF, JWH, RA and JAN; Funding acquisition, PSJB and JAN; Resources, RA, CF and JAN; Supervision, JAN.

## Acknowledgements

This work was supported by a Wellcome Senior Clinical Research Fellowship to JAN (102770/Z/13/Z and 215477/Z/19/Z), a Lister Institute Research Fellowship to JAN, and a Wellcome PhD Training Fellowship for Clinicians to PSJB (205252/Z/16/Z). Work in the CF laboratory was supported by the Medical Research Council (MRC_MC_UU_12022/6). This work was also supported by the NIHR Cambridge Biomedical Research Centre and Evelyn Trust. The Cambridge Institute for Medical Research is in receipt of a Wellcome Strategic Award (100140).

## Declaration of Interests

The authors declare no competing interests.

## Notes

### Competing Interest Statement

The authors have declared no competing interest.

