## Supplemental Files for "ABHD11 regulates 2-oxoglutarate abundance by protecting mitochondrial lipoylated proteins from lipid peroxidation damage"

Supplementary Information

**Figure S1. ABHD11 loss leads to HIF-1α accumulation and inhibition of 2-OG dependent dioxygenases in aerobic conditions.**

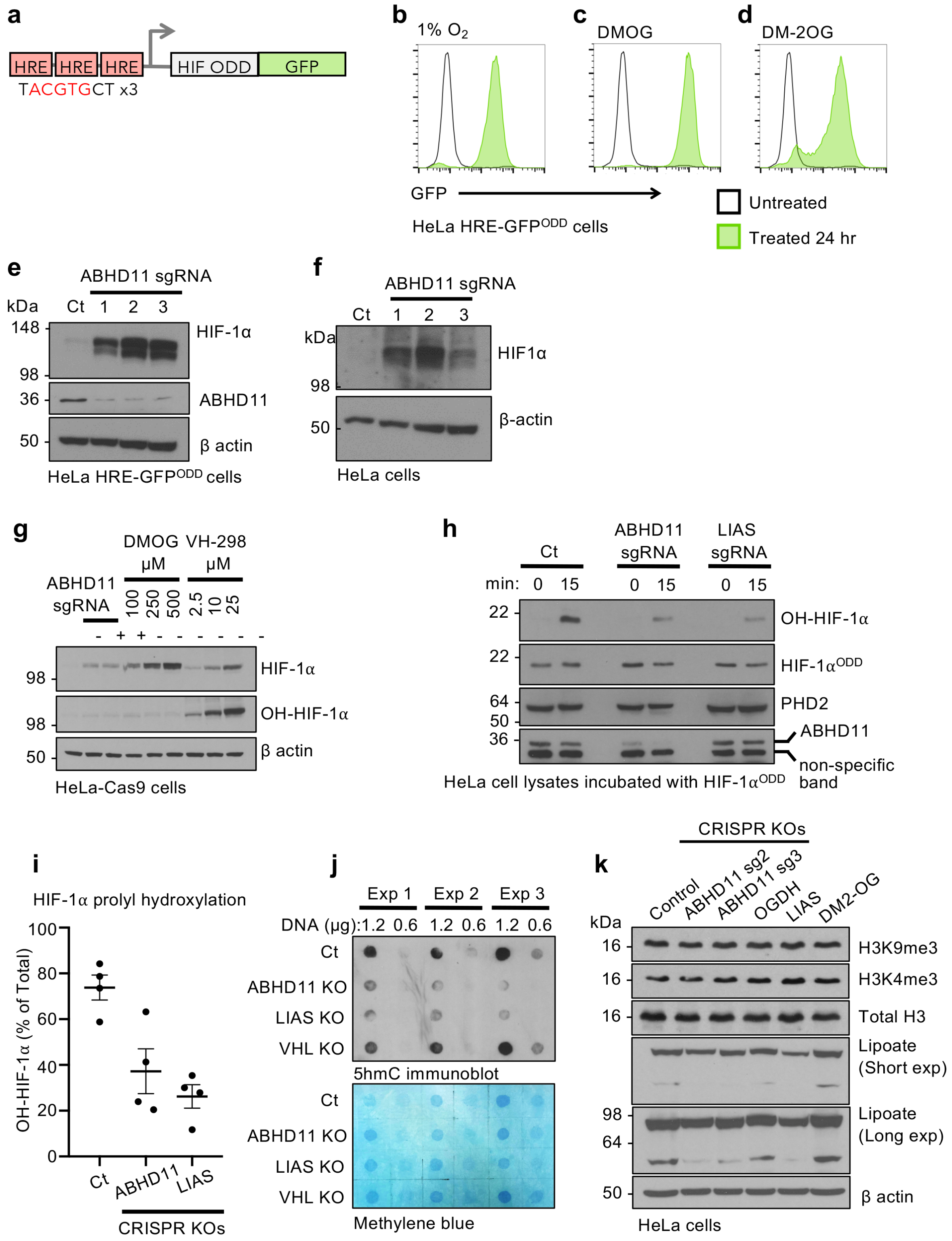

(**a**) Schematic of HRE-GFP$^{ODD}$ reporter. (**b-d**) HeLa HRE-GFP$^{ODD}$ cells respond to oxygen and metabolic inhibition. In 21% oxygen ($O_2$), GFP reporter levels are low (black line). Reporter cells incubated for 24 hours in 1% $O_2$ (**b**) or following treatment with 1 mM DMOG (**c**), or 4 mM dimethyl 2-OG (DM-2OG) (**d**) accumulate GFP. (**e, f**) ABHD11 depletion leads to HIF-1$\alpha$ accumulation in cells incubated in 21% oxygen. HeLa HRE-GFP$^{ODD}$ cells (**e**) or HeLa wildtype cells (**f**) expressing Cas9 were lentivirally transduced with up to 3 sgRNA targeting ABHD11 and HIF-1$\alpha$ levels measured by immunoblot. $\beta$-actin served as a loading control. (**g**) HIF-1$\alpha$ prolyl hydroxylation levels in HeLa cells following ABHD11 depletion, DMOG treatment or VHL inhibition. Control or ABHD11 deficient HeLa cells were generated as described. Hela cells were treated with indicated concentration of DMOG for 24 hr or the VHL inhibitor VH298 for 2 hours. Total HIF-1$\alpha$ and prolyl hydroxylated HIF-1$\alpha$ (OH-HIF-1$\alpha$) were measured by immunoblot. (**h, i**) *In vitro* prolyl hydroxylation of HIF-1$\alpha$. Recombinant HIF-1$\alpha$, encoding the C-terminal ODD region (aa 530 – 652), was incubated with cell extracts from control, ABHD11 deficient or LIAS deficient cells for 15 min at 37°C. Total HIF-1$\alpha^{ODD}$ and prolyl hydroxylated HIF-1$\alpha^{ODD}$ (OH-HIF-1$\alpha$) were measured by immunoblot (**h**) and quantified using ImageJ (**i**). *n=4, SEM.* (**j**) 5hmC levels are reduced in ABHD11 depleted cells. Genomic DNA was extracted from Hela control or mixed KO populations of ABHD11, LIAS or VHL, and 5hmC levels measured by immunoblot, compared to total DNA content measured by methylene blue stain (**Figure 1j**). (**k**) Effect of ABHD11 loss on histone methylation marks. Control, ABHD11, OGDH or LIAS deficient HeLa cells were generated as previously described. Wildtype Hela cells were also treated with DM2-OG (6mM 24hr) as indicated. H3K4me3, H3K9me3 and total H3 levels were measured by immunoblot. Short and long lipoate immunoblot exposures are shown.

**Figure S2. ABHD11 is a mitochondrial matrix hydrolase required for lipoylation of the OGDHc.**

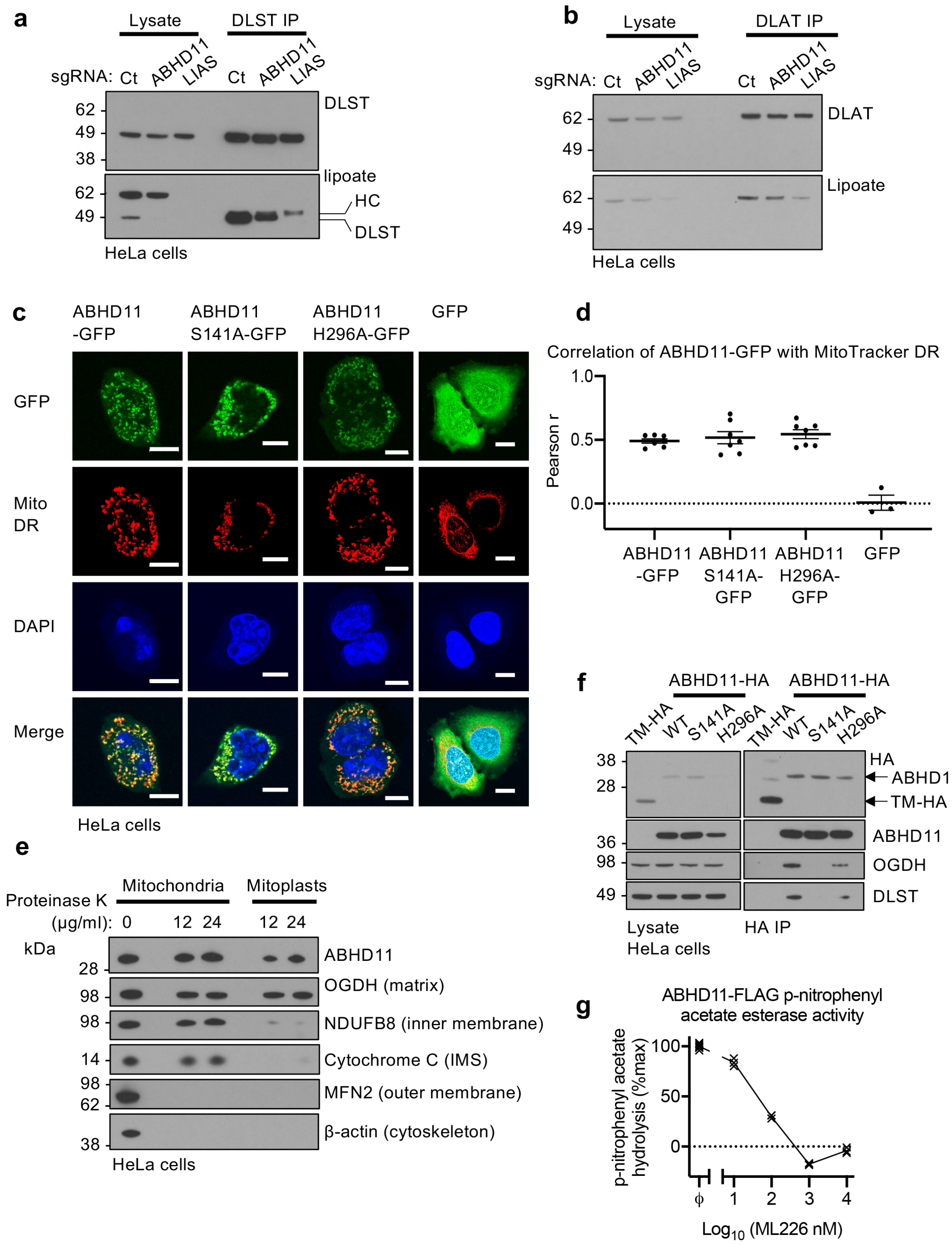

**(a)** Lipoylation of immunoprecipitated DLST. Control, ABHD11 mixed KO or LIAS mixed KO cells were lysed in 1% IGEPAL CA-630 and DLST immunoprecipitated. DLST and lipoylated DLST levels are shown. *HC = heavy chain.* (**b**) Lipoylation of immunoprecipitated DLAT. Control, ABHD11 mixed KO or LIAS mixed KO cells were lysed in 1% IGEPAL CA-630 and DLAT immunoprecipitated. DLAT and lipoylated DLAT levels are shown. (**c, d**) Confocal immunofluorescence microscopy of Hela cells stably expressing ABHD11-GFP, ABHD11 S141-GFP or ABHD11 H296A-GFP (**c**). MitoTracker Deep Red was used to visualise mitochondria. Pearson correlation coefficient was used to quantify ABHD11 colocalisation with mitochondria (**d**). *Scale=10$\mu$m*. (**e**) Mitochondrial protease protection assay. Mitochondria were extracted using the Qproteome Mitochondria Isolation Kit (Qiagen). Proteinase K was added to the final concentrations indicated, and incubation at 37°C for 30 minutes. Mitochondria or mitoplasts were lysed and analysed by immunoblot. (**f**) Immunoprecipitation of ABHD11-HA with endogenous OGDHc components. ABHD11-HA or the inactive mutants (S141A and H296A) were transduced into HeLa cells, lysed and immunoprecipitated using the HA tag. TMEM199, a membrane bound protein tagged with HA (TM-HA) was used as a control. (**g**) ML226 treatment in vitro. Purified wildtype ABHD11-FLAG was incubated with p-nitrophenyl acetate and hydrolysis measured by rate of increase in absorbance at 405 nm (37°C for 30 min), with addition of ML226 at the indicated concentration. Hydrolysis activity was subtracted from a background control without ABHD11-FLAG, and is normalised to the activity of ABHD11-FLAG with vehicle control. *$\phi$=vehicle control; n=3.*

**Figure S3. ABHD11 loss does not alter PDHc activity.**

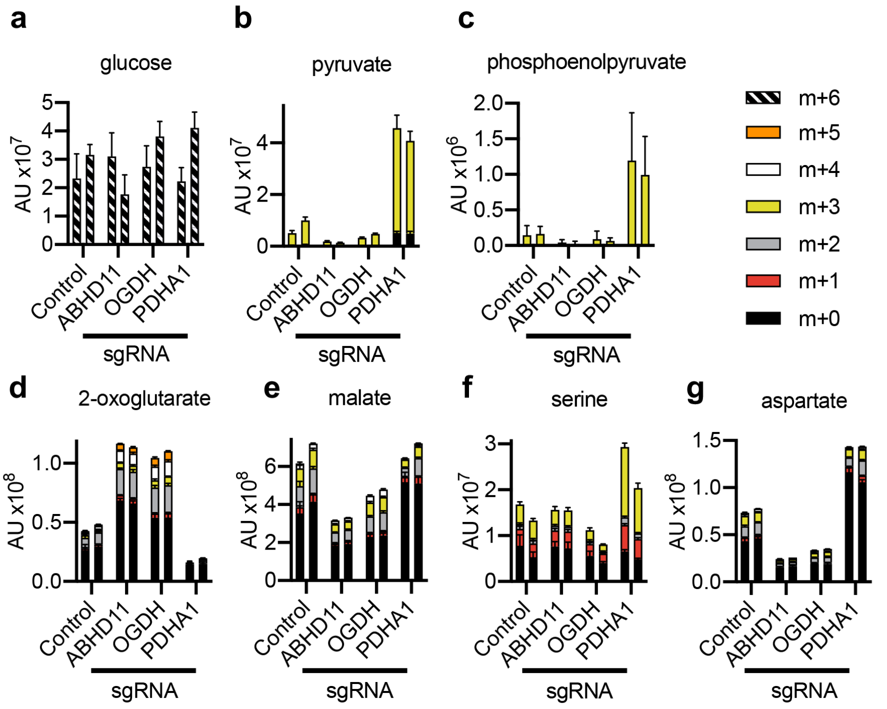

(**a-g**) Stable isotope tracing in HeLa cells compared to mixed CRISPR KO populations (sgRNA) of ABHD11, OGDH or pyruvate dehydrogenase E1 alpha subunit (PDHA1) incubated with [U-$^{13}$C6]-glucose. Isotopologues of pyruvate (**b**), and phosphoenolpyruvate (PEP) (**c**) confirm that PDHA1 loss impaired PDHc function, resulting in pyruvate and PEP accumulation. Isotopologues of 2-OG (**d**) and malate (**e**) confirm that ABHD11 and OGDH loss impairs the TCA cycle by decreased OGDHc activity, within increased 2-OG and decreased malate. Serine (**f**) and aspartate (**g**) increases following PDHA1 loss are consistent with decreased PDHc function. ABHD11 and OGDH loss do not increase serine or aspartate pools (**f, g**). Two biological replicates are shown with five technical repeats. The m+0 to m+5 isotopologues are indicated.

**Figure S4. Affinity purification of ABHD11-FLAG.**

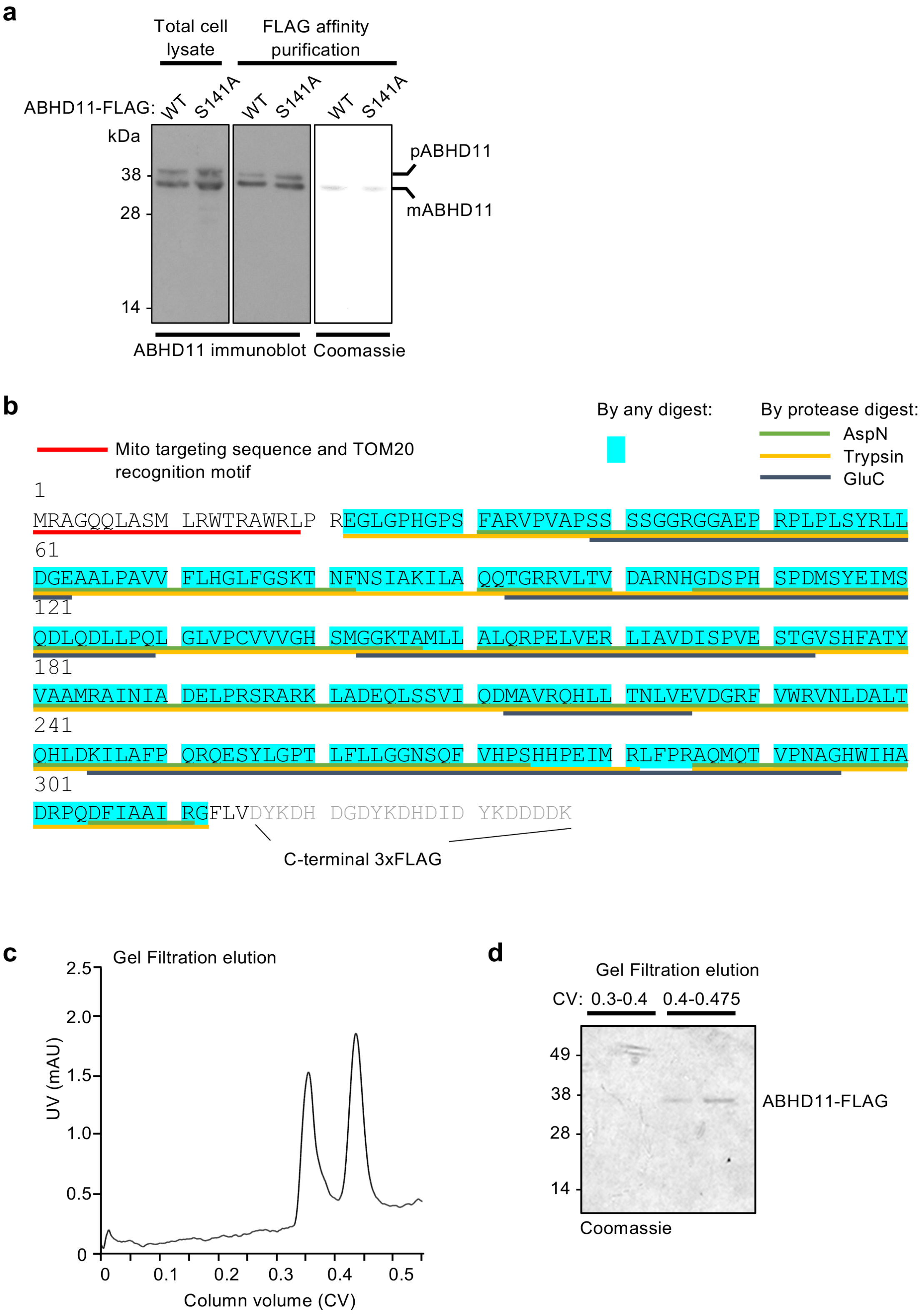

(**a**) Purification of wildtype or S141A mutant ABHD11 from human cells. HEK293T cells were transfected with wildtype of S141A ABHD11-FLAG. Cells were lysed after 48 hr (left panel) and ABHD11 affinity purified using anti-FLAG affinity beads and FLAG peptide elution (middle (immunoblot) and right panel (Coomassie staining)). The pre-cleaved (pABHD11) and mitochondrial cleaved (mABHD11) forms are indicated. (**b**) Mass spectrometry (MS) sequence analysis of affinity purified ABHD11-FLAG expressing HEK293T cells. Affinity purified ABHD11 was subjected to SDS-PAGE and analysed by MS using three different peptide digests (Trypsin, AspN or GluC). Complete peptide coverage was observed aside from the first 21 residues encoding the mitochondrial targeting sequence and TOM20 recognition motif. (**c, d**) Size exclusion chromatography of ABHD11-FLAG. Affinity purified ABHD11-FLAG was subjected to gel filtration using a Superdex 75 10/300 GL column. Two peaks were visualised, potentially corresponding to monomeric and dimeric forms. ABHD11-FLAG was only visualised in the second peak elution (**d**).

**Figure S5. ABHD11 prevents the formation of lipoyl adducts on the OGDHc**

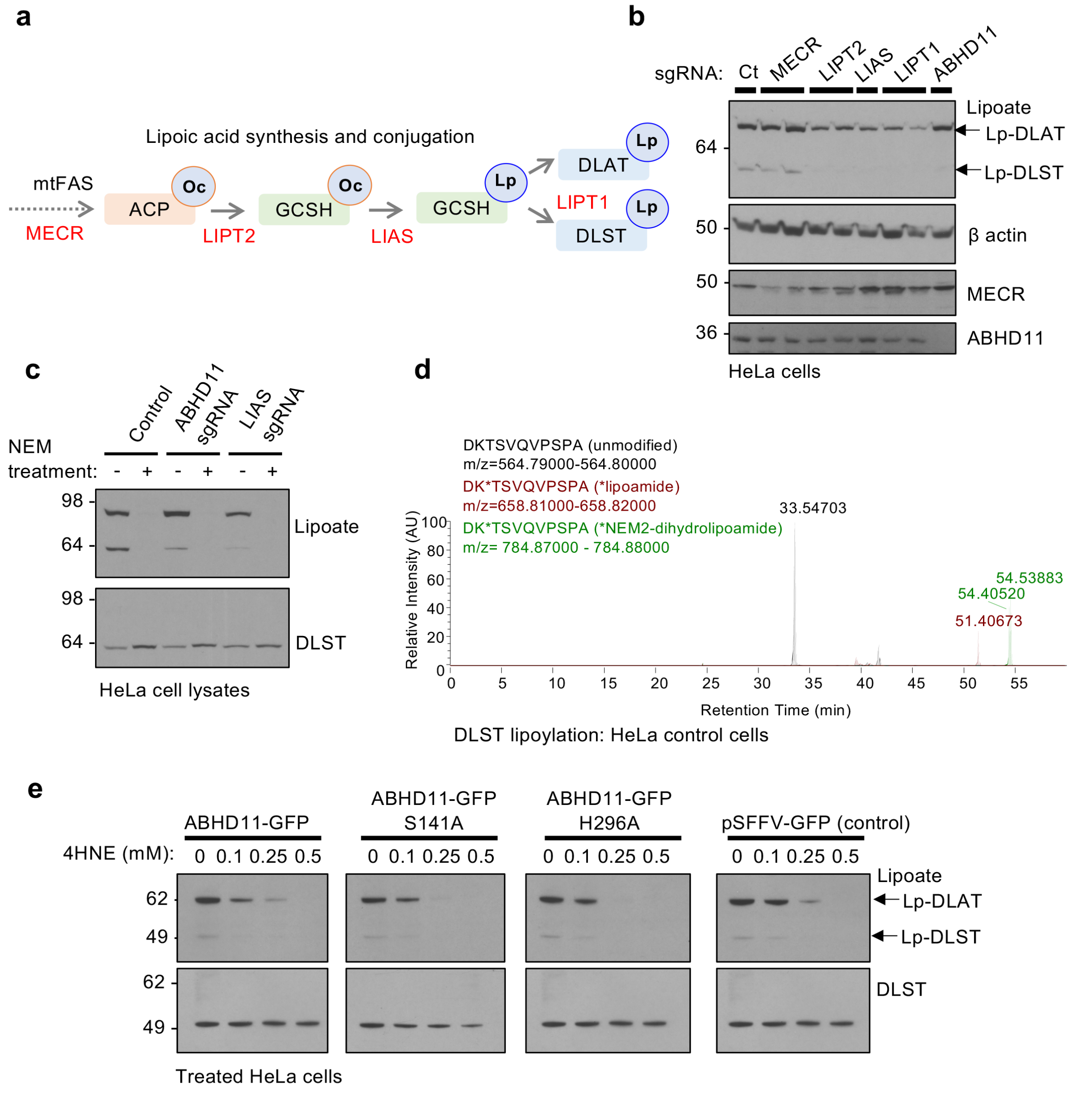

(**a**) Schematic of lipoic acid synthesis and conjugation pathway. Mitochondrial fatty acid synthesis provides the octanoylated (Oc) precursor for lipoic acid synthesis. MECR is upstream of octanoylated ACP, converting Trans-2-enoyl-ACP to acyl-ACP. LIPT2 catalyses the transfer of octanoate to GCSH. LIAS converts octanoate to lipoate. LIPT1 is thought to be the major lipoyl transferase allowing conjugation to conserved lysine residues in 2-oxoacid dehydrogenase subunits such as DLAT and DLST. (**b**) Depletion of lipoic acid synthesis and conjugation components in HeLa cells. Mixed HeLa KO populations of MECR, LIPT2, LIPT1, LIAS and ABHD11 were generated and immunoblotted for lipoylated proteins. β-actin served as a loading control. (**c**) Effect of N-ethyl maleimide (NEM) treatment on protein lipoylation detected by immunoblot. Control or ABHD11 deficient HeLa cells were lysed in a HEPES buffer, and free thiols in HeLa cells were blocked with the addition of 10mM NEM (4°C, 1 hour). Immunoblot of lipoylated proteins confirmed efficient blocking of free thiols. (**d**) Representative chromatograms of DLST peptide encoding the lipoylated region from HeLa cells, demonstrating that the m/z separation of the differently modified DLST peptide (DK*TSVQVPSPA). (**e**) Overexpression of ABHD11 catalytic inactive mutants renders cells more susceptible to lipoyl adduct formation. HeLa cells expressing ABHD11-GFP, ABHD11 S141A-GFP, ABHD11 H296A-GFP or a GFP only lentivirus (pSFFV-GFP) were treated with 4-HNE at the concentrations indicated, and lipoylation levels measured using the anti-lipoate antibody. Loss of lipoylation represents lipoyl adduct formation.

**Table S1.**

| **Reagents and Antibodies** | Source | Identifier |
|---|---|---|
| **Antibodies** | | |
| Rabbit polyclonal anti-5hmC | Active Motif | Cat#39770; RRID: AB_10013602 |
| Rabbit polyclonal anti-ABHD11 | Signalway | Cat#34366 |
| Mouse monoclonal anti-β actin | Sigma | Cat#A2228; RRID: AB_476697 |
| Mouse monoclonal anti-Cytochrome C | Abcam | Cat#ab110325; RRID: AB_10864775 |
| Rabbit polyclonal anti-DLD | GeneTex | Cat#GTX101245; RRID: AB_1240715 |
| Mouse monoclonal anti-DLAT | Cell Signalling | Cat#12362S; RRID: AB_2797893 |
| Mouse monoclonal anti-DLST (used for IP) | Abcam | Cat#ab110306; RRID: AB_10862702 |
| Rabbit monoclonal anti-DLST (used for immunoblot) | Cell Signalling | Cat#11954; RRID: AB_2732907 |
| Mouse monoclonal anti-GFP | Roche | Cat#11814460001; RRID: AB_390913 |
| Rat monoclonal anti-HA (used for immunoblot) | Roche | Cat#11867423001; RRID: AB_390918 |
| Mouse monoclonal anti-HIF-1α | BD Biosciences | Cat#610959; RRID: AB_398272 |
| Rabbit monoclonal anti-Hydroxy-HIF-1α | Cell Signalling | Cat#3434; RRID: RRID:AB_2116958 |
| Rabbit polyclonal anti-Lipoic acid | Calbiochem | Cat#437695; RRID: AB_212120 |
| Mouse monoclonal anti-MFN2 | Abcam | Cat#ab56889; RRID: AB_2142629 |
| Mouse monoclonal anti-NDUFB8 | Abcam | Cat#ab110242; RRID: AB_10859122 |
| Rabbit polyclonal anti-OGDH | Atlas Antibodies | Cat#HPA020347; RRID: AB_1854773 |
| Rabbit polyclonal anti-PHD2 (EGLN1) | Novus | Cat#NB100-137; RRID: AB_10003054 |
| **Reagents** | | |
| Antimycin A | Alfa Aesar | Cat#J63522; CAS: 1397-94-0s |
| Dimethyloxalylglycine | Cayman Chemical | Cat#71210; CAS: 89464-63-1 |

| | | |
|---|---|---|
| FCCP | Cayman Chemical | Cat#15218; CAS: 370-86-5 |
| Sodium oxamate | Sigma | Cat# O2751; CAS: 565-73-1 |
| GSK-2837808A | Tocris | CAS: 1445879-21-9 |
| ML226 | Cayman Chemical | Cat#25681; CAS: 2055172-43-3 |
| Oligomycin A | Cayman Chemical | Cat#11342; CAS: 579-13-5 |
| Rotenone | Sigma | Cat#R8875; CAS: 83-79-4 |
| VH298 | A gift from Alessio Ciulli | |
| 3xFLAG Peptide | Sigma | Cat#F4799 |
| His-HIF-1$\alpha^{ODD}$ (aa530-652) | Burr et al, 2016 | |

**Table S2**

| | |
|---|---|
| **CRISPR sgRNA oligonucleotide sequences** | |
| ABHD11 sgRNA 1 | TGCTGTAGATATCAGCCCAG |
| ABHD11 sgRNA 2 | AAGATCTTGGCCCAGCAGAC |
| ABHD11 sgRNA 3 | GCTGTGGCCAACGACGACGC |
| ABHD11 sgRNA 4 | GCAGAAGGTCCTGCAGGTCC |
| LIAS sgRNA 1 | TTAGGTTAAGACTACCTCCA |
| LIPT1 sgRNA 1 | ATGCCTACCAATTACAACAG |
| LIPT1 sgRNA 2 | GTAGCCTGCACATCCAGCTG |
| LIPT2 sgRNA 1 | CAGGCGCACCAACCGAACGG |
| LIPT2 sgRNA 2 | TATACGGCCGGGCTGCGCGG |
| MECR sgRNA 1 | GCAGGGATTGACACCCAGGG |
| MECR sgRNA 2 | GAAGTCCATCAACATCCTGT |
| OGDH sgRNA 1 | CTGCTCTTACCTCCAGCCGA |
| OGDH sgRNA 2 | TTCCTGTCCCCCGATGAAAG |
| PDHA1 sgRNA 1 | GATGCAGACTGTACGCCGAA |
| PDHA1 sgRNA 2 | AGGATGGGCTCAAATACTAC |
| PHD2 sgRNA 1 | ATGCCGTGCTTGTTCATGCA |
| VHL sgRNA 1 | GTGCCATCTCTCAATGTTGA |
| **Custom screen primers (TKO)** | |
| TKO Inner PCR forward | AATGATACGGCGACCACCGAGATCTACA CTCTCTTGTGGAAAGGACGAGGTACCG |
| TKO custom sequencing primer | ACACTCTCTTGTGGAAAGGACGAGGTACCG |
| **NEBuilder HiFi PCR primers to clone ABHD11 into pHRSIN lentiviral vector** | |
| Forward | CAGTCCTCCGACAGACTGAGTCGCCCGGGGG GGATCCGCCACCATGcgagccggccaacagcttgcaa |
| Reverse | CTTGCATGCCTGCAGGTCGACTCTAGAGTCGC GGCCGCttagaccaggaagcctcggatggcagc |
| **NEBuilder HiFi PCR primer to clone ABHD11 with C-terminal GFP tag into pHRSIN lentiviral vector** | |
| Reverse (for ABHD11) | CTTGCTCACgaccaggaagcctcggatggc |
| Forward (GFP) | gcttcctggtcGTGAGCAAGGGCGAGGAGCTG |
| Reverse (GFP) | GTCGACTCTAGAGTCGCGGCCGCttaCTTGTA CAGCTCGTCCATGCCGAG |
| **NEBuilder HiFi PCR primer to clone ABHD11 with C-terminal HA tag** | |
| Reverse | GTCGACTCTAGAGTCGCGGCCGCttaCT TGTACAGCTCGTCCATGCCGAG |

| | |
|---|---|
| **NEBuilder HiFi PCR primers to create silent mutations in ABHD11 sgRNA site 2 (mutations capitalised)** | |
| Forward | caaAatACtCgcAcaAcaAacaggccgtagggtg |
| Reverse | gtTtgTtgTgcGaGTatTttggcgatggagttgaagttag |
| **NEBuilder HiFi PCR primers to create S141A active site mutation in ABHD11** | |
| Forward | cacGCGatgggaggaaagacag |
| Reverse | catCGCgtggccaacgacgac |
| **NEBuilder HiFi PCR primers to create H296A active site mutation in ABHD11** | |
| Forward | ccgaacgctggcGCc |
| Reverse | cagcgtggatccagGCg |
| **Gibson Assembly PCR primers to clone ABHD11 into pCEFL 3xFLAG mCherry vector** | |
| Forward | GGAATTGGCGAAGCTTGGTACCGAGCTCGG ATCCGCCACCATGcgagccggccaacagcttgcaa |
| Reverse | CCGTCATGGTCTTTGTAGTCAGCCCGCTCG AGCGGCCGCCCgaccaggaagcctcggatggcagc |

List of sgRNA sequences used, and PCR primers used for genome-wide screen amplification and sequencing, and cloning of ABHD11 expression vectors.
